# An integrative field and modelling study of the bottom-up, top-down and indirect effects of native species on invasive insects and their biological control : the case of the worldwide chestnut tree pest, *Dryocosmus kuriphilus*, in the French Eastern Pyrenees

**DOI:** 10.1101/2025.09.21.677588

**Authors:** Jean-Loup Zitoun, Raphaël Rousseau, Sébastien Gourbière

**Affiliations:** UMR5096 ‘Laboratoire Génome et Développement des Plantes’, Université de Perpignan Via Domitia, Perpignan, France; School of Life Sciences, University of Sussex, Falmer, Brighton, United Kingdom

## Abstract

Insect pests invading an ecosystem typically face various forces regulating their population growth and spread. While the resources they feed on and their natural enemies have ‘bottom-up’ and ‘top-down’ effects on insect invasion success, other native species, more distant in the local trophic network, can have indirect effects on the ecosystem’s susceptibility to invasion. Resolving the mechanisms underlying these naturally occurring indirect interactions and their impact on invasion dynamics is a key challenge toward risk assessment and environmental management. In this contribution, we investigated how indirect interactions with several native species contribute to variations in *Dryocosmus kuriphilus* infestation level in 24 natural chestnut tree populations of the French Eastern Pyrenees. This invasive gall-forming hymenopteran parasites cultivated and wild *Castanea sativa* stands, and its hymenopteran parasitoid, *Torymus sinensis*, is world widely used as a control agent. We combined ecological, molecular and statistical approaches to quantify the effects of *Quercus pubescens*, *Fagus sylvatica* and the parasitic fungus *Cryphonectria parasitica* on the pest oviposition rate (*effect i*), its host detection capacity (*effect ii*), the production of native parasitoids (*effect iii*) and in providing alternative hosts for the control agent (*effect iv*). The integration of these effects into a specifically designed *D. kuriphilus – T. sinensis* dynamical model, provided clear evidence of the quantitative impacts of *Q. pubescens* and *C. parasitica* on the invasion potential of the pest and its biological control.

## Introduction

Over the last two centuries, the frequency of biological invasions has dramatically increased worldwide (1). Invasive species represent a major threat for biodiversity in natural ecosystems (2) as they are the main driver of plants and animals extinctions (3), involved in hybridization with native species (4), spreading pathogens worldwide (5) and perturbing trophic networks (6). Damages caused by insect pests on natural biological resources and attempts to limit them stand as key causes of economic losses (7), with an annual cost reaching US$26.8 billion worldwide (8). When invading an ecosystem, any pest faces natural ‘bottom-up’ and ‘top-down’ controls associated with the resources they feed on and their natural enemies, as well as indirect sources of regulation through more indirect interactions with other native species present in the local environment (e.g. 9-11). Resolving the mechanisms underlying such direct and indirect interactions and their impact on invasion dynamics represents one of the major challenges to assess ecosystem’s susceptibility to invasion and set-up efficient control strategies.

Bottom-up regulation factors depend on the resources used by an invasive species to grow its population and are typically related to its feeding requirements. The distribution, genetic and physiological characteristics of such resources can indeed impact the establishment and spreading success of a pest (12–13). On the other hand, top-down regulation factors correspond to the predators, parasites and pathogens that can directly affect the survival or reproductive rates of an invasive species and therefore reduce its local growth (14–15). Although feeding resources and natural enemies typically are the main local determinants of an invasion through their direct impact on the alien species (e.g. 9), other species, more distant in the trophic network, can have indirect effects on their spread and control. According to the host (plant) visual apparency and the semiochemical redundancy hypothesis (16–17), the presence of species phylogenetically close to the resource used by the invasive pest could induce a so-called ‘dilution effect’ by lowering the ability of the pest population to detect its resource (18–19). Furthermore, variations in the physiological characteristics of the resource induced by pathogen infection have also been shown to contribute in modulating the spread of invasive pests (20). Indirect interactions with native species can also affect the efficacy of biological control strategies targeting invasive pests by leading to predation pressure on the control agent (21), inter-specific competition for the resource (22–23) or by providing alternative hosts or refuges (24).

The indirect contributions of such species remain broadly overlooked as invasion ecology is usually focused on the delicate partitioning of the role of bottom-up or top-down factors (9,25–26). The integration of empirical knowledge of the (direct) bottom-up and top-down determinants of invasion into dynamic models provides an efficient approach to assess their relative impacts on the spread of exogenous species in newly invaded environments (e.g. 27-28). The analysis of such modelling, further accounting for the indirect effects of native species, would allow for quantitative predictions about their contributions to typical features of the invasion and its control, such as the rate of spread of the invasive (29–30) and the dynamical outcomes of the interaction with its control agents (31–32).

We intended to develop such an approach to improve our understanding of the determinants of the spread of the invasive *Dryocosmus kuriphilus*, the most virulent pest of chestnut trees (*Castanea sativa*) orchards and forests worldwide (33). This cynipid, originated from China, has spread across Asia (34), North-America (35) and Europe (36–37), where its galling activity on chestnut trees inhibits the development of the shoots and ultimately lead to reduction of up to 80% of fruit production (38) and of up to 60% of the annual tree-ring increment (39). Its rapid and successful invasion history is attributed to a very effective parthenogenetic reproduction, the lack of natural enemies in the invaded ranges and its long range dispersal assisted by human-mediated transport of infested plant material (40). The control agent, *Torymus sinensis*, has been introduced worldwide to restrain the spread of the pest and reduce its impact on both cultivated and (semi-)natural chestnut tree populations. This univoltine and semelparous hymenoptera of the *Torymidae* family showed successful establishment and effective control rates in Japan, North America and Europe (40). The impact of bottom-up and top-down regulation factors on the *D. kuriphilus* spread and its biological control in the natural chestnut tree forests of the French Eastern Pyrenees have recently been investigated using ‘eco-genomic’ models (28,41). While these integrative approaches have shown that the chestnut tree frequency and genetic susceptibility are the main natural determinants of *D. kuriphilus* invasion potential that can also impact the success of its biological control, the potential effect of indirect interactions with native species were not assessed.

In this contribution, we aim at producing quantitative insights into four main hypothetical mechanisms by which native species could have an indirect impact on the spread of *D. kuriphilus* and its control agent, *T. sinensis,* in the French Eastern Pyrenees. First, *Quercus pubescens* and *Fagus sylvatica*, two local tree species related to *C. sativa*, could lower the ability of *D. kuriphilus* to detect its chestnut tree hosts through the perturbation of odor and visual clues (17–18). Second, infection of chestnut trees by the fungal parasite *Cryphonectria parasitica* have been shown to increase the host detection of *D. kuriphilus* because of the enhanced release of phenolics compounds induced by canker damages (42). Third, local tree species, i.e. *Q. pubescens* and *F. sylvatica*, that are hosts of native gall-forming parasites, could increase the abundance of native parasitoids populations (43) and/or, fourth, the abundance of the introduced parasitoids used as control agent, *T. sinensis*, since the latter has been shown to be able to exploit such alternative plant hosts (44). To assess the impact of these four hypothetical mechanisms, we set-up a three years ecological field study to estimate levels of *D. kuriphilus* and *C. parasitica* infestation and the tree community structure in 24 sampling sites distributed across 8 localities representative of the chestnut tree populations of the French Eastern Pyrenees. We concomitantly identified the parasitoids, both native and introduced, infesting galls formed by native gall-forming parasites on *Q. pubescens* and *F. sylvatica*. The above effects were then integrated into a dynamical model of the *D. kuriphilus - T. sinensis* interaction to provide further quantitative insights into the impacts of these native species on the pest spread and its biological control.

## Material and Methods

### Dryocosmus kuriphilus life-cycle and its bottom-up and top-down control

Univoltine and semelparous females of *D. kuriphilus* emerge from galls in June-July and fly to lay their asexually produced eggs in chestnut tree buds during their 1-7 days short life-span (45). Oviposition can trigger a local immune response in chestnut trees buds that prevent the development of the deposited eggs (46). This host resistance mechanism has been shown to exhibit substantial variations between chestnut tree strains, ranging from totally-resistant individuals to highly susceptible ones (47). Eggs that survive the host response and intrinsic developmental failure, hatch within a month and develop into first instar larvae that stay in dormant stage to overwinter. In spring, they emerge from dormancy and develop into subsequent larval stage to induce the formation of galls during the chestnut tree buds burst, where they eventually become pupae and adults that will produce the next and non-overlapping generation of eggs. Although *D. kuriphilus* galls have been shown to be targeted and infected by local fungus (48) and hymenopteran parasitoids (28,40,49), the additional mortality they induce on the pest larvae remains generally lower than 10% and does not allow for a natural regulation of its invasion neither in the French Eastern Pyrenees (28,41) nor elsewhere (50–51).

### Tree community structure and chestnut tree infestation by D. kuriphilus and C. parasitica

The forest tree community structure was estimated in each of the 24 study sites located in the 8 localities selected across the chestnut tree populations of the French Eastern Pyrenees. We sampled 1000m² of forest in each site and identified the taxonomic status of each tree to the species level, which provided estimates of the frequency of *C. sativa*, *Q. pubescens* and *F. sylvatica.* The level of infestation of the local chestnut tree populations by *D. kuriphilus* were estimated from 2019 to 2021 by using two typical measures of parasitism. The infestation rate, defined as the mean number of galls per leaf, was estimated on 5 geo-located chestnut trees per sampling site and with a sampling effort of 250 to 500 leaves per tree. The prevalence rate, defined as the proportion of chestnut trees infested by at least one gall, was estimated on 50 chestnut trees per site. The prevalence of *C. parasitica* infection, defined as the proportion of chestnut trees presenting chestnut blight symptoms, was assessed in 2021 on the same 50 chestnut trees in each of the 24 sampling sites. The existence of heterogeneity between sites or localities in the prevalences and rates of chestnut trees infestation by *D. kuriphilus and C. parasitica* was tested by Pearson’s chi-squared tests with Yates’s correction for continuity. Exact confidence intervals for both measures of infestation were calculated from binomial distributions set according to the corresponding sampling efforts.

### General Linear Mixed Models Analyses (GLMMs) and statistical tests

In order to test the effects of *Q. pubescens, F. sylvatica* and *C. parasitica* on *D. kuriphilus* ability to infest its chestnut tree hosts, we conducted 2 independent GLMMs analyses. Those allowed to establish the relationships between (i) prevalence and (ii) rate of *D. kuriphilus* infestation observed among the 24 sampling sites, and the four following explanatory variables; frequencies of *C. sativa* (among the whole tree community), *Q. pubescens* and *F. sylvatica* (among the non-host tree community) and the prevalence of *C. parasitica infection,* that were all included as continuous fixed-effects. Since the sampling year had a significant impact on both infestation indexes (Prevalence: Estimate = −1.7243, p = 9.05 10^-11^, Infestation: Estimate = −1.3674, p = 3.77 10^-8^), we accounted for such a temporal structure by following the procedure recommended by Zuur et al. (52). We applied model simplification by starting with the highest order interaction between the above explanatory variables and by sequentially removing non-significant variable and interaction predictors. All analyses were conducted in R 4.4.2 (53) using the glmmPQL function from the MASS package. We further tested the possibility of a positive association between *D. kuriphilus* and *C. parasitica* infections of chestnut trees using a Pearson’s chi-squared tests with Yates’s correction for continuity.

### Collection of the native gall forming parasites and their parasitoids on Q. pubescens and F. sylvatica

The prevalences and rates of infestation of *Q. pubescens* (by the gall-forming parasites *Andricus kollari* and *Andricus dentimitratus)* and of *F. sylvatica* (by the gall-forming parasite *Mikiola fagi*) were estimated using the same protocol as for the infestation of chestnut trees by *D. kuriphilus* (see above). The infestation rate, defined as the mean number of galls per leaf, was estimated on the same 5 geo-located chestnut trees per sampling site and with a similar sampling effort of 250 to 500 leaves per tree. The prevalence rate, defined as the proportion of chestnut trees infested by at least one gall, was estimated on 50 chestnut trees per site. To identify which native or introduced parasitoid species could develop in the galls formed on *Q. pubescens and F. sylvatica,* we dissected 56 galls of *A. kollari*, 152 galls of *A. dentimitratus* galls and 263 galls of *M. fagi*. These dissections allowed for the collection of 50, 149 and 201 larvae, respectively, that were then divided into 7 pools for DNA extraction (see below). Larvae found in galls of *A. kollari* (AK) and *A. dentimitratus* (AD) were distributed in 3 pools corresponding to the ‘Massane’ locality, where 52% of the galls (and 64% of the larvae) were found, and in 2 pools for the 6 other sampling stations (Céret, Saint-Laurent, Laroque and Prats-de-mollo, Arles-sur-Tech, Llauro) where the remaining pubescent oaks galls were collected. The larvae found in galls of *M. fagi* (MF) were separated between the two localities where they were collected (i.e., ‘Bastide’ and ‘Massane’).

### Molecular identification of the parasitoids found in galls of Q. pubescens and F. sylvatica parasites

DNA extractions were performed for each pool of larvae using the E.Z.N.A Tissue DNA kit (Omega BIO-TEK). A Polymerase Chain Reaction (PCR) was performed to amplify the ribosomal Internal Transcribed Spacers 2 (ITS2), with primers obtained from Viviani et al. (54). PCRs were performed in 35 μL reaction volume, containing ∼ 20 ng DNA template, 0.1 μM of each dNTP, 0.04 μM of each primer and 0.7 U Phusion High-Fidelity DNA polymerase (FINNZYMES OY, Espoo, Finland) in 7 μL 5X manufacturer’s buffer plus 21,35 μL sterile distilled water. The thermal profile of the PCR was as follows: initial denaturation at 98°C for 30 seconds, followed by 16 to 22 cycles (depending on the sample) of denaturation at 98°C for 10 seconds, locus-specific annealing at 52,7°C for 30 seconds, elongation at 72°C for 18s, and a final elongation at 72°C for 10 minutes. Libraries were generated using Nextera index and Q5 high fidelity DNA polymerase (New England Biolabs). PCR products were normalized with SequalPrep plates (Thermofischer). Paired-end sequencing was performed on a MiSeq system (Illumina) using 2×300bp v3 chemistry. Sequences obtained from the 7 pools of samples were processed with the FROGS pipeline (55) available on the Genotool Bioinfo galaxy server (56). Clustering swarm step was made with aggregation distance of 1. Only OTU’s with at least 50 sequences were conserved and affiliated based on the NCBI database.

### Integration of native species interferences into a D. kuriphilus – T. sinensis dynamic model

To provide further quantitative insights into how native species could influence the pest spread and its biological control, we integrated the empirically supported effects of *Q. pubescens* and *C. parasitica* into a dynamical model of *D. kuriphilus* - *T. sinensis* interactions that we had previously developed (28). This modelling is based on the seminal Nicholson-Bailey model (57) that is widely used in theoretical ecology to describe host-parasitoids population dynamics and which was earlier adapted to account for the effect of top-down and bottom-up determinants on the invasive pest population (Figure 1). Here, we further expanded this initial modelling to account for the i) reduction of *D. kuriphilus* oviposition associated with the presence of *Q. pubescens*, the ii) facilitative effect of chestnut blight symptoms on *D. kuriphilus* host detection, and the productions of iii) native parasitoids and iv) introduced *T. sinensis* in galls formed on *Q. pubescen*s.

**Figure 1.**
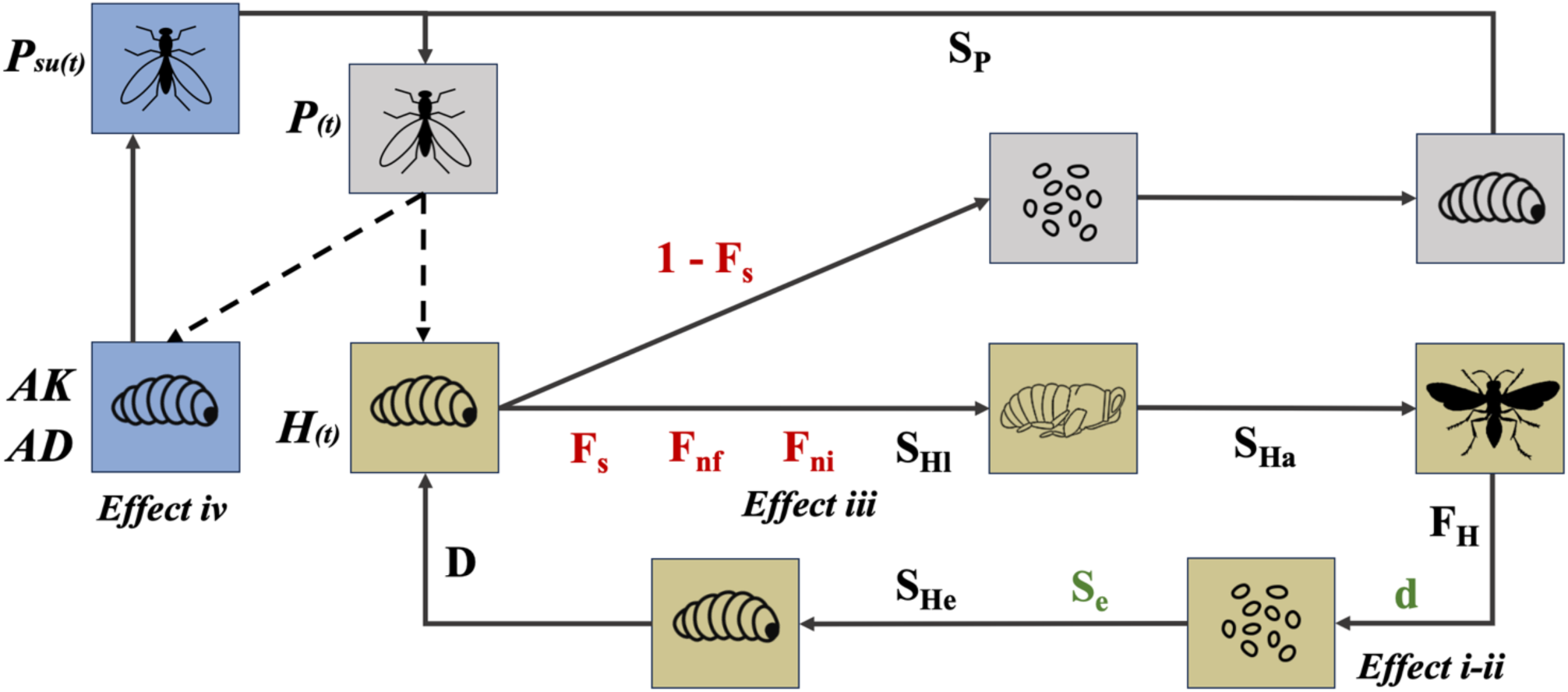
Dynamical model of the *D. kuriphilus – T. sinensis* interaction accounting for interferences with *Q. pubescens* and *C. parasitica.* The lower part of the graph depicts the different stages of *D. kuriphilus* life-cycle (yellow) with the effects of the native hyperparasitic insects and fungi (i.e., the ‘top-down’ control appearing in red) and the impacts of the tree community and host genetic diversity (i.e., the ‘bottom-up’ control shown in green). The upper part of the graph describes the life-cycle of the introduced control agent, *T. sinensis* (grey), that is able to lay its eggs in both *D. kuriphilus* larvae (found in galls formed on chestnut trees) and larvae of *A. kollari* and *A. dentimitratus* (found in galls formed on *Q. pubescens*). All parameters are defined in the main text and their estimates are provided in SM2 to SM4.

In the modelling shown in Figure 1, the *D. kuriphilus* (host) larvae (H(t)) are split into those exploited by *T. sinensis* adults (P_s_(t)) to lay their eggs, and those escaping the control agent. According to the low ability of *T. sinensis* to fly towards the galls of *D. kuriphilus* (58), the fraction of *D. kuriphilus* larvae escaping the parasitoid (F_s_) was defined by assuming a random dispersal of *T. sinensis* within its searching area (*a*_s_) and using a Poisson distribution. Under such an assumption, the probability of being parasited by no egg at all, i.e. to escape parasitism by *T. sinensis*, corresponds to F_s_(H(t), P_s_(t) P_c_) = e^-a^_s_^,P^_c_^,P(t)^, where p_c_ stands for the frequency of chestnut trees in the forest environment.

The larvae of *D. kuriphilus* escaping *T. sinensis* face additional challenges by native hyperparasite fungi and insects that exert a constant infective pressure, which larvae can escape with probabilities F_ni_ and F_nf_, respectively. The production of native parasitoids in galls formed on *Q. pubescen*s (*effect iii*) was introduced through an increase in parameter F_ni_ with the frequency of pubescent oaks, as further specified below. Larvae infected by none of these hyperparasites develop into pupae at rate S_Hl_ and moult into adults with probability S_Ha_. After emergence, the *D. kuriphilus* females produce an average of F_H_ eggs per individual, and a fraction d of those eggs are deposited in chestnut tree buds. The effects of *Q. pubescens* and *C. parasitica* on *D. kuriphilus* host detection and oviposition (*effects i* and *ii*) were accounted for by using a typical host-choice function to model d with respect to the frequencies of healthy chestnut trees, blight infected chestnut trees, pubescent oaks, and the preferences of *D. kuriphilus’* for each of these trees, as further specified below. To develop into larvae and enter in dormancy to overwinter, the deposited eggs must then survive to a tree hypersensitive response and intrinsic causes of mortality during development, which they do with probabilities S_e_ and S_He_, with the former depending on the observed distribution of *C. sativa* susceptibility, as estimated in Zitoun et al. (28). The description of *D. kuriphilus* life-cycle, was completed by modelling the density dependent survival of dormant larvae associated with the competition for space during chambers formation. Such a typical ‘contest’ competition was described using a function proposed by Brännström and Sumpter (59): 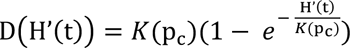, where H’(t) stands for the number of larvae entering dormancy and K(p_#_) for the maximal amount of *D. kuriphilus* larvae that can be sustained in a stand, which was set to increase linearly with the proportion p_c_ of chestnut-trees, as estimated in Zitoun et al. (28).

The complementary part of *D. kuriphilus* larvae, i.e. the fraction 1-F_s_ not escaping parasitism by *T. sinensis,* is set to die as the hyperparasite larvae feed on them to become, with probability S_P_, the next generation of adults that will intend to lay their eggs into *D. kuriphilus, A. kollari* and *A. dentimitratus* larvae. To account for the production of *T. sinensis* associated with the oviposition in larvae of *A. kollari* and *A. dentimitratus*, whose galls are formed on *Q. pubescen*s (*effect iv*), we assumed a constant number of larvae available in such galls, and described their infection by *T. sinensis* according to specific searching efficiency for these hosts, as further specified below.

The modelling of the population dynamics of *D. kuriphilus* and *T. sinensis* accounting for both the direct ‘bottom-up’ and ‘top-down’ control, as well as the indirect effects of native tree and fungal species (*i-iv*) can then be described by the following set of non-linear difference equations:

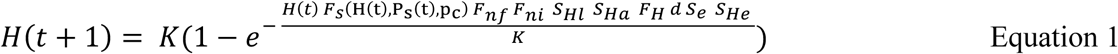

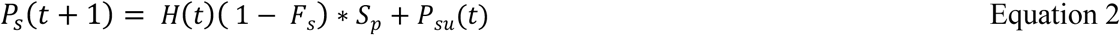

where *P*_su_(*t*) stands for the amount of *T. sinensis* adults produced in *Q. pubescen*s galls in year t, and is defined according to equation 5 (see below).

### Impact of Q. pubescens and C. parasitica on D. kuriphilus’ host detection and oviposition

To integrate the effects induced by pubescent oaks and chestnut blight infection on *D. kuriphilus* host detection and oviposition (*effects i* and *ii*), the fraction of eggs deposited on chestnut tree (d) was formally linked, through a typical host-choice function, with the frequencies of healthy chestnut trees (p_hc_), blight-infected chestnut trees (p_ic_), pubescent oaks (p_u_), and the preferences of *D. kuriphilus* for each of these trees (a_hc_, a_ic_ and a_u_) and other non-host trees (a_n_):

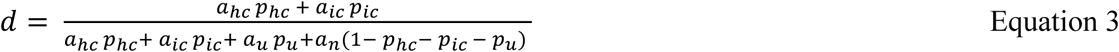

The pest ability to detect healthy (a_hc_), blight-infected (a_ic_) chestnut trees and other non-host tree species (a_n_) were estimated from data of Germinara et al. (42). The ability of *D. kuriphilus* to detect *Q. pubescens* was assumed to be similar to its ability to detect healthy chestnut trees as both species are closely related. All estimates (shown in SM2) were derived by considering a detection capacity of 1 in the absence of preference, and the pest’s capacity to detect a tree category as linearly proportional to the observed preference for this category.

### Impact of Q. pubescens on the production of native parasitoids

To account for the indirect effect of *Q. pubescens* associated with the production of native parasitoids (*effect iii*), we assumed that the percentage of *D. kuriphilus* larvae hyperparasited by such native parasitoids is merely proportional to the density of pubescent oaks, as those correspond to their natural ecological niche. The probability to escape such a parasitism can then be written:

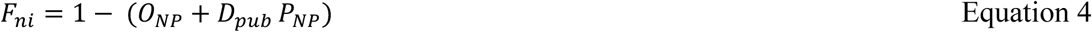

where O_NP_ and P_NP_ correspond to the y-intercept and slope of the linear relationship between the emergence rate of native parasitoids and the density of pubescent oaks (D_pub_). This linear relationship was estimated from the rates of *D. kuriphilus* hyperparasitism and densities of pubescent oaks measured across our 24 sampling sites (SM2).

### Integration of the production of T. sinensis on Q. pubescens

To account for the production of *T. sinensis* associated with the hyperparasitism of *A. kollari* and *A. dentimitratus* larvae in galls formed on *Q. pubescen*s *(effect iv),* we modelled the rate of such hyperparasitism by assuming a random dispersal of the parasitoid within its searching area. We then used a similar Poisson distribution as considered to model the rate of p hyperparasitism of *D. kuriphilus* larvae (see above), which we defined according to specific searching efficiencies of *T. sinensis* for these alternative host. The amount of *T. sinensis* adults produced on *Q. pubescen*s can then be written as the sum of the production on both native parasitic species:

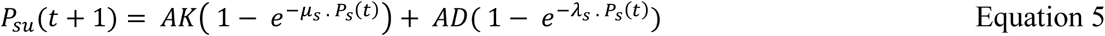

where μ_s_ and λ_s_ represent the *T. sinensis* searching efficiency for *A. kollari* and *A. dentimitratus* larvae, whose point estimates and 95% confidence intervals are presented in SM3, and where AK and AD stand for the abundances of *A. kollari* and *A. dentimitratus* larvae, respectively.

### Model predictions on D. kuriphilus invasion potential and its biological control

All model predictions were made numerically using an implementation of our dynamical model (Equations 1 to 5) in R 4.4.2 (53). We first aimed to look at the changes in the invasive capacity of *D. kuriphilus* induced by the i) dilution of *D. kuriphilus* oviposition by *Q. pubescens*, the ii) facilitative impact of *C. parasitica* on *D. kuriphilus* host detection, and the iii) production of native parasitoids in galls formed on *Q. pubescen*s. To do so, we accounted for one of these three effects (i to iii) at a time, and estimated *D. kuriphilus’s* invasion potential (measured by its R_0_) in 15 forest environments by setting the frequency of chestnut trees to its minimal (0.16), average (0.60) or maximal (0.91) values estimated across our 24 sampling sites, and by letting the frequency of *Q. pubescens* (among non-chestnut trees) or the prevalence of *C. parasitica* infection to take on five different values (0; 0.25; 0.5; 0.75; 1). In each of these 15 environmental settings, we calculated *R_0_* by using the estimates of *D. kuriphilus* life-history parameters and hyperparasitism by native fungi species (provided in SM4) and by accounting for the uncertainty in our estimates of i) the *D. kuriphilus* preferences (for *C. parasitica* infected and non-infected chestnut trees and for pubescent oaks) and of ii) the intercept and slope of the linear relationship between *Q. pubescens* densities and chestnut tree galls infestation by native parasitoid species. To account for such uncertainty we randomly sampled 200 values of these three quantities using their confidence intervals (described in SM3), which provided a distribution of *R_0_* values represented by boxplots in Figure 4. These variations were shown in Figure 4A-C, and further expressed per additional percent in the presence of native species (i.e. in the frequency of *Q. pubescens* or the prevalence of *C. parasitica*) introduced in the modelled chestnut tree forest considered in Figure 4D-F.

Second, we aimed at using our modelling to predict how much the biological control of the invasion by the introduced agent, *T. sinensis,* could be increased by its use of the galls of *A. kollari* and *A. dentimitratus* found on the native oak species *Q. pubescens.* We then used our modelling to predict the proportions of *A. kollari* and *A. dentimitratus* oak galls infested by *T. sinensis.* These predictions were made by running simulations for the minimum, average and maximum value of the 95% confidence interval of the searching areas of *T. sinensis* for *A. kollari* and *A. dentimitratus* (provided in SM4) and for different frequencies of chestnut tree ranging from 0 to 1. When the *D. kuriphilus – T. sinensis* interaction dynamics led to sustained oscillations, which typically occurs when the chestnut tree frequency is larger than 70% (28), we further recorded the minimal and maximal values of these rates of infestation, which all appear in Figure 5. Finally, we predicted the impact of *Q. pubescens* and *C. parasitica* (*effect i-iv*) on the biological control efficacy (SM5) by running simulations with and without the control agent, and by estimating the level of biological control as the ratio between the abundance of *D. kuriphilus* found in both circumstances.

## Results

### Significant and uneven decreases of the chestnut trees infestation by *D. kuriphilus*

The studied chestnut tree populations of the French Eastern Pyrenees were all found infested by *D. kuriphilus* in 2019, and the infestation level decreased across the entire area in both 2020 and 2021, with heterogeneous reduction rates among the 24 sampling sites (SM1) and the 8 localities (Figure 2).

**Figure 2.**
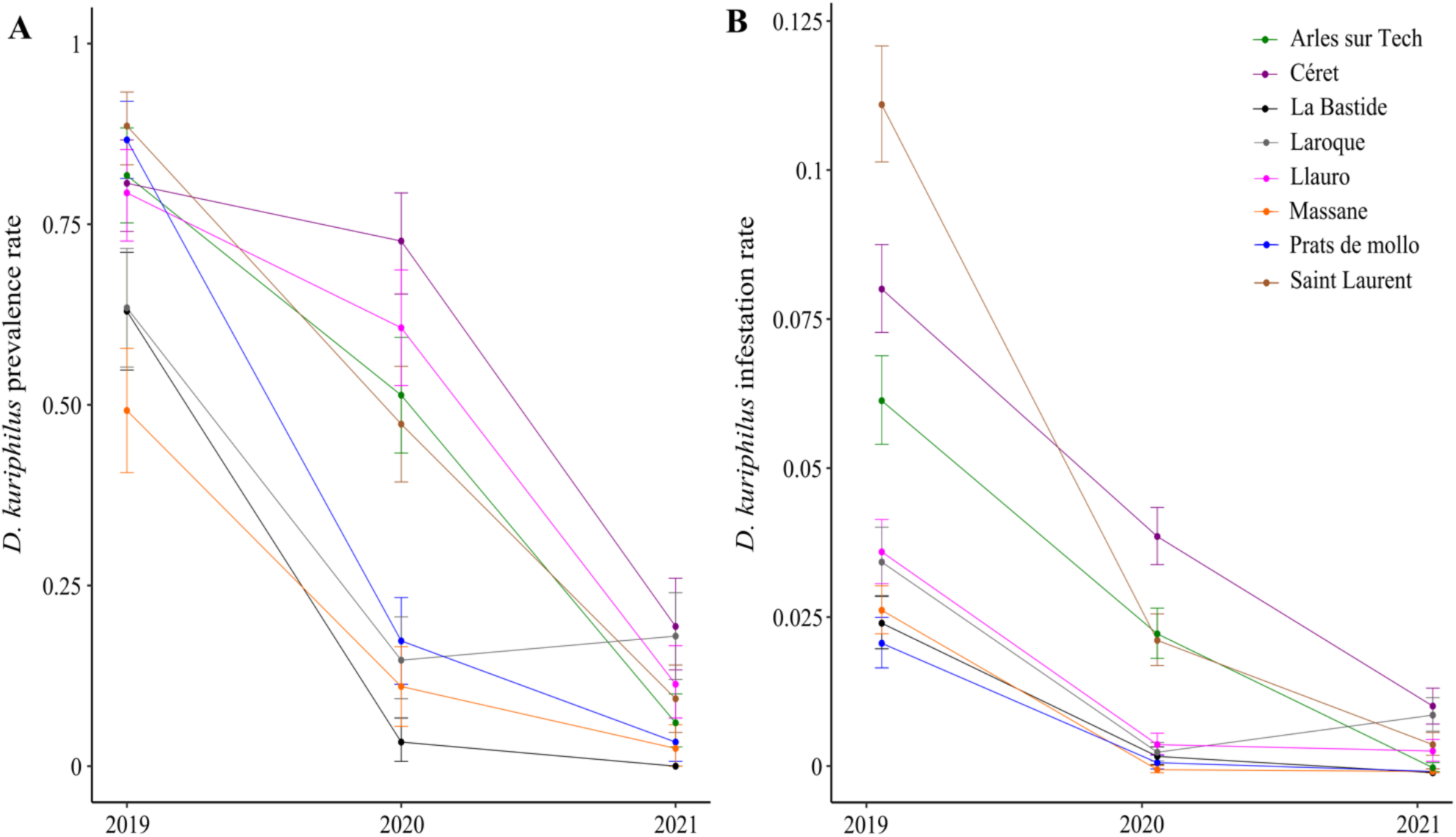
Infestation levels of chestnut tree populations by *D. kuriphilus* in the French Eastern Pyrenees. Measures of the prevalence (A) and rate (B) of infestation made in 2019, 2020 and 2021 and in the 8 studied localities (named in the upper right panel) appear with their 95% confidence intervals.

In 2019, 75% of individual chestnut trees were infested with an average of 0.05 galls per leaf, representing a parasitic burden of 1 gall every 20 leaves. The overall proportion of infested trees then dropped to 34% with an average of 0.013 galls per leaf in 2020, and to 9% with an average of 0.004 galls per leaf in 2021 (Figure 2, SM1). The prevalence of infestation (proportion of infested trees) showed spatial heterogeneity between sites (χ^2^ = 271.45, df = 71, p < 2.2 10^-16^) and between stations (χ^2^ = 160.96, df = 23, p < 2.2 10^-16^), and similar trends were observed for the rate of infestation (average number of galls per leaf) between sites (χ^2^ = 1056.6, df = 71, p < 2.2 10^-16^) and stations (χ^2^ = 682.3, df = 23, p < 2.2 10^-16^).

The decreases in prevalence and infestation rates observed between 2019 and 2020 represent overall reductions of 55% and 74% in these two infestation measures, while further drops of 74% and 69% were observed between 2020 and 2021, leading to mean reductions of 88% and 92% over these two years period (Figure 2). Although a decrease in both prevalence and infestation rate was observed in almost all sampling sites between 2019 and 2021 (SM1), significant variations in the amplitude of changes were observed in prevalence between stations (χ^2^ = 16.815, df = 7, p = 0.01863) and sites (χ^2^ = 53.618, df = 23, p = 0.000302) and in the rate of infestation (χ^2^ = 261.46, df = 7, p < 2.2 10^-16^, χ^2^ = 466.29, df = 23, p < 2.2 10^-16^). Notably, 6 sampling sites, 3 of which located in the ‘Laroque’ locality, even showed an increase in infestation between 2020 and 2021.

Overall, *D. kuriphilus* was found to be widely spread across the chestnut tree populations of the French Eastern Pyrenees with a strong spatial heterogeneity in prevalence and infestation rates observed at both locality and site scales, and with both measures of infestation pointing toward a decrease in *D. kuriphilus* abundance in almost all localities sampled between 2019 and 2021.

### Chestnut tree-dominated forests suffer heterogeneous levels of canker infestation

Chestnut trees were found predominant in 19 of the 24 sampling sites where their overall frequency ranges from 15.8% to 90.8% (Figure 3, SM1). Pubescent oak was present in 17 sites with frequency varying between 0.8% and 20.7%, while beech was recorded only in 3 sites with frequency ranging from 3% to 56%. The local estimates of the tree community structure in sampling sites were consistent with the largest scale description of vegetal formations found in national inventories, as sites located in ‘Chestnut tree stand’ typically showed higher chestnut tree frequencies as compared to those located in ‘deciduous’ and ‘coniferous and deciduous forests’ (0.51 ± 0.01) (Figure 3). Canker symptoms were observed on chestnut trees in 19 sites (Figure 3, SM1) and showed significant spatial heterogeneity in the infestation levels (χ^2^ = 115.32, df = 23, p = 2.871 10^-14^). Overall, 17.6% of (207/1177) chestnut trees were found infected by *C. parasitica,* with such prevalence rate varying between 2% and 52% between sites.

**Figure 3.**
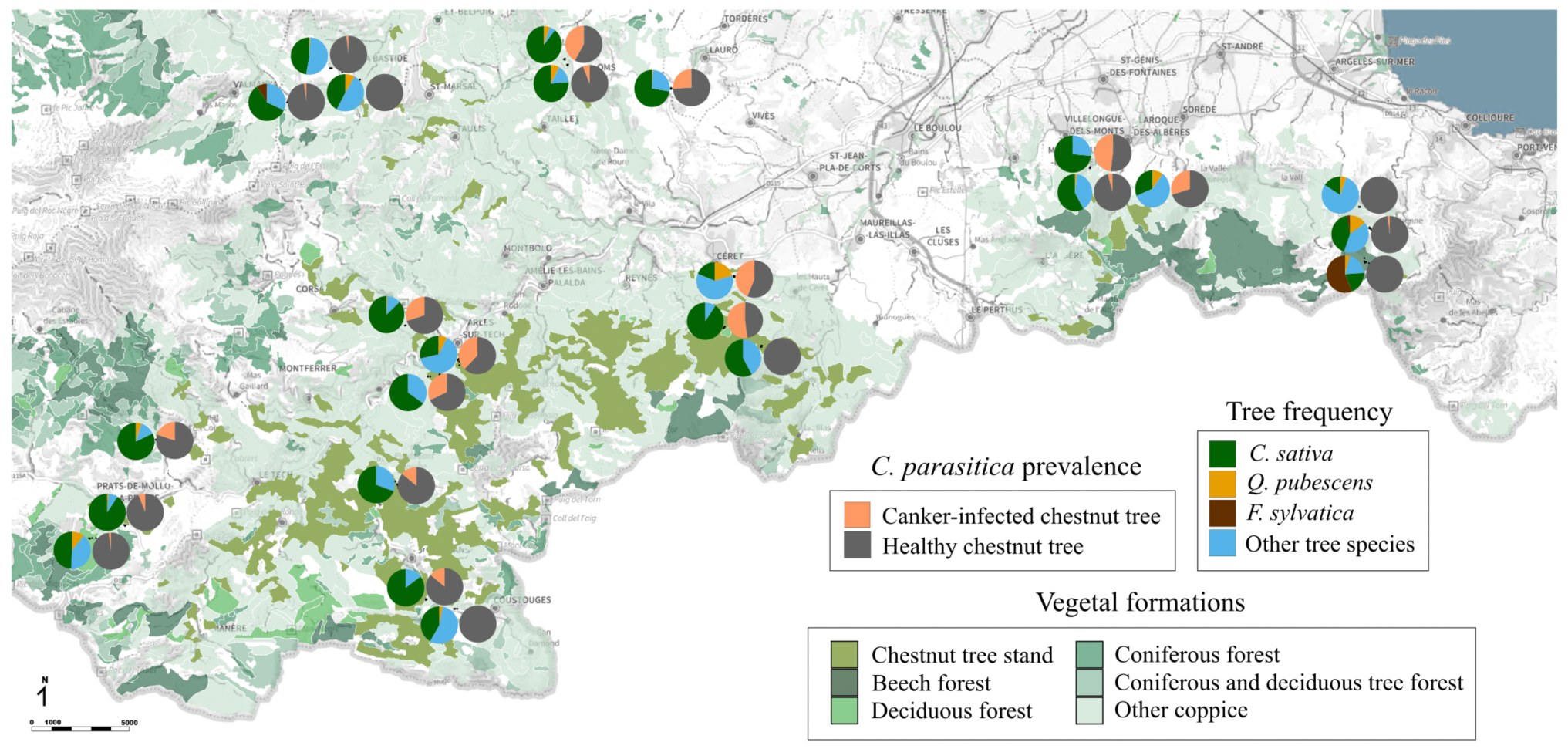
Map of the vegetal formations in French Eastern Pyrenees with tree species frequencies and *C. parasitica* prevalence measured in the 24 sampling sites. Pie charts describe the frequencies of *Q. pubescens, F. sylvatica, C. sativa* and *C. parasitica* prevalence estimated in the 24 sampling sites. The different vegetal formations containing chestnut trees are represented on the map with different shade of green specified in the lower right legend panel.

We then intended to look at the impact of the native tree and parasitic fungal species, *Q. pubescens, F. sylvatica* and *C. parasitica,* to the spatial and temporal variations observed in the infestation of the chestnut tree populations by the invasive pest *D. Kuriphilus* (i.e. the effects i and ii).

### Pubescent oaks dilute *D. kuriphilus* oviposition while canker symptoms facilitate its hosts detection

The generalized linear mixed model (GLMM) regression analysis revealed that the observed variations in *D. kuriphilus* infestation rates between sites are positively correlated to the chestnut tree frequency, and negatively correlated to the frequency of pubescent oak among the non-host trees (Table 1A). These findings support the hypothesis that the release of odor clues by this phylogenetically close tree species (*Q. pubescens*) can perturbate dispersing insects and create a dilution effect lowering the pest’s oviposition rate. Meanwhile, the prevalence of *D. kuriphilus* infestation was positively correlated with the chestnut tree frequency and with the canker prevalence observed in each site (Table 1B). At the individual tree level, a higher number of chestnut trees were found infected by both *D. kuriphilus* and *C. parasitica* than expected under the assumption of independent parasitism (Table 1C) (χ^2^ = 57.972, df = 1, p = 2.7 10^-14^). These observations are consistent with the hypothesis that the detection of canker infected trees by *D. kuriphilus* is facilitated, presumably by the enhanced release of phenolics compounds.

**Table 1.** Statistical analysis of native species interferences with *D. kuriphilus* infestation levels. GLMM analysis of the correlation between canker prevalence, chestnut tree, pubescent oak and beeches frequency with infestation (A) and prevalence (B) rate at the site level. (C) Contingency table of the number of chestnut tree showing *D. kuriphilus* galls and/or canker symptoms. Numbers appearing in grey correspond to the abundances expected under the null hypothesis of independent infestation of chestnut tree hosts by *D. kuriphilus* and *C. parasitica*.

| Fixed effects | Estimate | Standard error | df | P |
| --- | --- | --- | --- | --- |
| **A** Response variable : *D. kuriphilus* infestation rate |  |  |  |  |
| Intercept | -4.755097 | 0.6966306 | 67 | 0 |
| Chestnut tree frequency | 1.368448 | 0.4554777 | 67 | 0.0037 |
| Pubescent oak frequency (among non-host trees) | -3.977703 | 0.9906581 | 67 | 0.0002 |
| **B** Response variable : *D. kuriphilus* prevalence rate |  |  |  |  |
| Intercept | -2.506713 | 0.9800511 | 67 | 0.128 |
| Chestnut tree frequency | 2.629348 | 0.6340386 | 67 | 0.0001 |
| Canker prevalence | 1.698257 | 0.8205657 | 67 | 0.0423 |

**Table 1. Statistical analysis of native species interferences with *D.**
| **C** | *D. kuriphilus* (-) | *D. kuriphilus* (+) |  |
| --- | --- | --- | --- |
| *C. parasitica* (-) | 676 (627.99) | 294 (342.01) | 970 |
| *C. parasitica* (+) | 86 (134.01) | 121 (72.99) | 207 |
|  | 762 | 415 |  |

The presence of gall-forming tree species, *Q. pubescens* and *F. sylvatica*, was also thought to have an impact on the *D. kuriphilus* populations by allowing for the production of native parasitoids (*effect iii*) and/or by providing an additional ecological niche for the control agent, *T. sinensis* (*effect iv*). We then characterized the parasitoid communities associated with galls formed on these two tree species to assess the presence of native parasitoids species *D. kuriphilus* galls, and of the control agent *T. sinensis*.

### *Q. pubescens* galls contain both native parasitoids infesting *D. kuriphilus* galls and *T. sinensis*

We dissected 471 galls collected on *Q. pubescens* (208) and *F. sylvatica* (263) in order to identify the different species of native or introduced parasitoids able to hyperparasite the larvae of the local hymenoptera, *A. kollari* and *A. dentimitratus*, and diptera, *M. fagi*, that induce the formation of those galls on pubescent oaks and beeches, respectively. *A. kollari* galls were found on pubescent oaks in 21 of our sampling sites, with 34% of individual trees being infested and an average rate of 0.004 galls per leaf. *A. dentimitratus* galls were observed in 18 sites, with 18% of pubescent oaks infested and an average rate of 0.011 galls per leaf. *M. fagi* galls were observed on beeches in all 3 stations where the tree species was present. All individual trees were then found infested with an average rate of 0.102 galls per leaf (SM1). From the 471 galls formed by the three local parasite species, we obtained 400 larvae that were split into 7 pools for molecular taxonomic identification (Table 2).

**Table 2.** Parasite and parasitoid species identified in *Q. pubescens* and *F. sylvatica* galls. Molecular taxonomic identification of the larvae contained in galls of *A. kollari* (AK), *A. dentimitratus* (AD) and *M. fagi* (MF) and found in the geographic pools denoted (1) = Céret, Saint-Laurent and Laroque; (2) = Prats de mollo, Arles sur Tech and Llauro; (MA) = Massane, and (BA) = La Bastide. n stands for the number of larvae in each subset.

| Species | AKAD-1<br>n = 39 | AKAD-2<br>n = 33 | AKAD-MA<br>n = 66 | AD-MA<br>n = 45 | AK-MA<br>n = 16 | MF-BA<br>n = 97 | MF-MA<br>n = 104 |
| --- | --- | --- | --- | --- | --- | --- | --- |
| *Andricus caputmedusae* | X | X | X | X |  |  |  |
| *Andricus kollari* | X | X | X |  |  |  |  |
| *Andricus quercustozae* | X | X |  | X |  |  |  |
| *Andricus sp.* | X | X | X |  |  |  |  |
| *Eulophidae sp* |  |  |  |  |  | X | X |
| *Eupelmus sp.* |  |  |  |  |  | X | X |
| *Eurytoma rosae* |  | X |  | X |  |  |  |
| *Eurytoma sp.* | X |  | X |  |  |  |  |
| *Megastigmus dorsalis* |  |  | X |  |  |  |  |
| *Megastigmus stigmatizans* | X | X | X | X | X |  |  |
| *Nasiona sp.* | X |  |  |  |  |  |  |
| *Synergus sp.* |  | X |  | X |  |  |  |
| *Torymus auratus* |  | X |  |  |  |  |  |
| *Torymus flavipes* |  |  | X |  |  |  |  |
| *Torymus sinensis* | X | X | X |  | X |  |  |
| *Torymus sp.* |  |  |  |  |  | X |  |

Galls of *A. kollari* and *A. dentimitratus* were found infested by at least 3 hymenopteran species known to infest *D. kuriphilus* larvae (*Torymus auratus, Torymus flavipes, T. sinensis*). Although native parasitoids (*T. auratus, T. flavipes)* were expected to be observed on their endemic hosts, the finding of the introduced *T. sinensis* in 4 out of 5 DNA pools originated from galls collected on *Q. pubescens* sharply contrasts with its description as a specialist parasitoid of *D. kuriphilus*. While the DNA of *Torymus sp.*, *Eulophidae sp.* and *Eupelmus sp.* were found in galls (of *M. fagi*) collected on beeches, the taxonomic resolution remained insufficient to establish that the corresponding species could potentially infest *D. kuriphilus* larvae. This analysis of the parasitoid community associated with *Q. pubescens* and *F. sylvatica* therefore provides evidence that galls found on pubescent oaks harbour native and introduced parasitoids able to infect *D. kuriphilus*. Furthermore, native parasitoids emergence rates from chestnut tree galls were shown positively correlated to pubescent oak abundances in the 24 sites (Spearman’s rank rho = 0.464, p = 0.022). These results supports the hypothesis that, beyond its dilution effect on oviposition, *Q. pubescens* presence could contribute to the regulation of *D. kuriphilus* invasion by increasing native parasitoid production (*effect iii*) and/or providing a new ecological niche for the control agent *T. sinensis* (*effect iv*). To provide further quantitative insights into the impact of *Q. pubescens* and *C. parasitica* on the *D. kuriphilus* invasion potential an control, we used our dynamical model of *D. kuriphilus – T. sinensis* interactions that accounts for their empirically supported effects (*i-iv*).

### Q. pubescens and C. parasitica induce significant variations of D. kuriphilus’ R_0_

We first assessed how the effects *i* to *iii* impact the growth rate (*R_0_*) of the pest population in forests with low, medium or high chestnut tree frequencies (Figure 4). The reduction of the oviposition rate of *D. kuriphilus* associated with the presence of *Q. pubescens* (*effect i*) lead to a systematic decrease of *R_0_* proportional to the frequency of *Q. pubescens* (Figure 4A). In a forest made of 60% of chestnut trees, i.e. the average value observed across our sampling stations, *D. kuriphilus’ R_0_* is reduced from 19.9 to 16.6 (−16%) when the frequency of pubescent oaks (among the remaining trees) is risen from 0 to 100%. As expected, the intensity of this dilution of *D. kuriphilus* oviposition was accentuated when considering the lowest frequency of chestnut trees measured in our sampling sites, i.e. 16%, as the frequency of *Q. pubescens* in the stand was then allowed to reach higher values up to 84%. The *D. kuriphilus R_0_* then decreased by up to 35%, falling from 6.8 to 4.4. Meanwhile, the pest’ *R_0_* was only reduced by 3.8% in forest with the maximal frequency of chestnut trees observed in our study, i.e. 91%.

**Figure 4.**
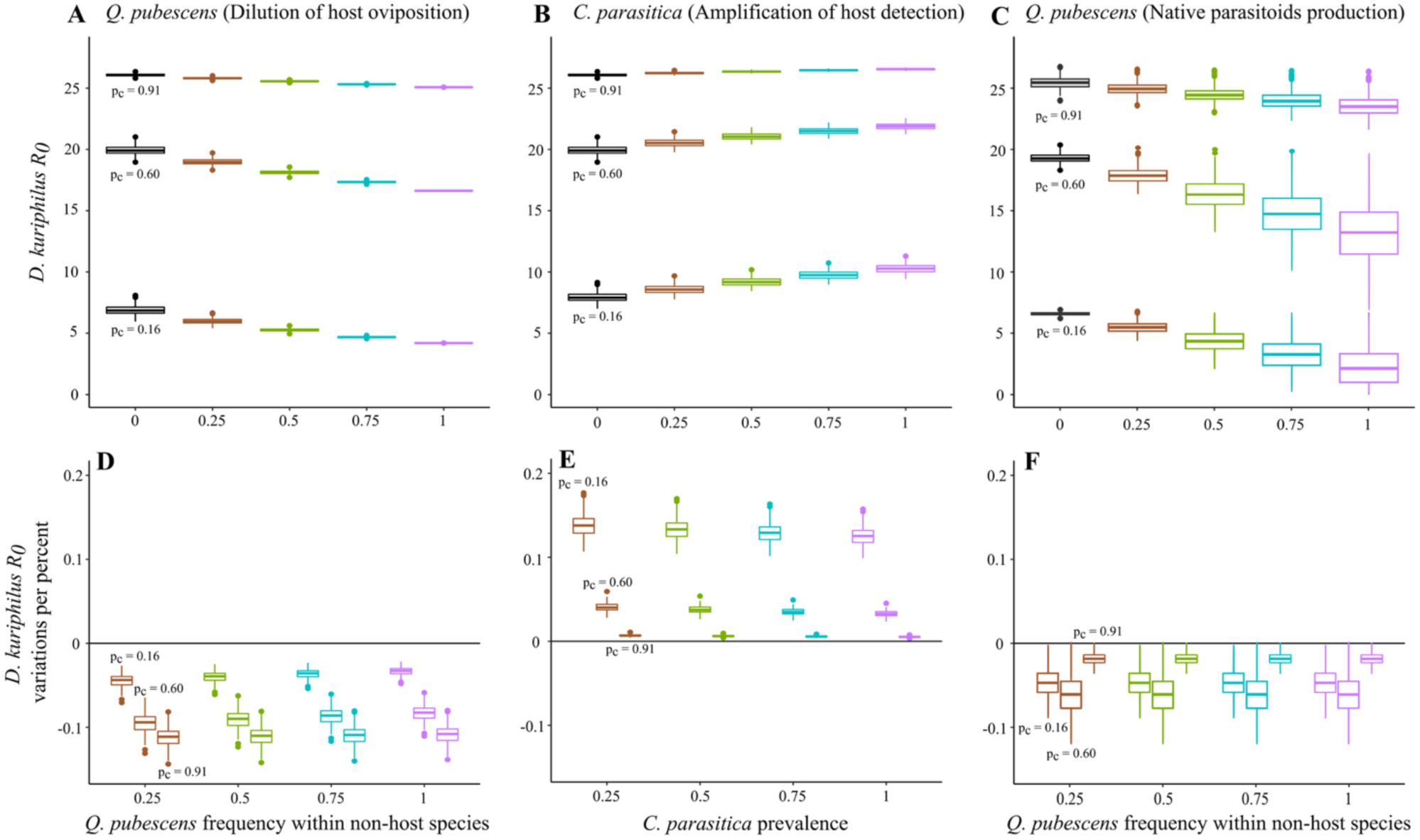
Variations of *D. kuriphilus R_0_* with the effects of *Q. pubescens* or *C. parasitica* in forests with low, medium and high abundances of chestnut trees. The variations of R_0_ are shown with respect to the effect of *Q. pubescens* (A) and *C. parasitica* (B) on *D. kuriphilus* oviposition and detection of chestnut trees, and according to the impact of *Q. pubescens* on the production of native parasitoids (C). The absolute variations of *R_0_* predicted in A-C were then transformed into a variation of *R_0_* per percent of change in the frequencies of native species (D,E,F). In each specific condition, the variability in the estimate of *R_0_* (calculated from the confidence intervals around the mean value of all its defining parameters, see ‘Material and Method’) was represented by a boxplot.

The amplification of *D. kuriphilus* host detection associated with chestnut tree infection by *C. parasitica* (*effect ii*) contributed to increase the pest’ *R_0_* proportionally to canker prevalence levels (Figure 4B). In forest with an intermediate frequency of chestnut trees, this amplification effect increased the value of *R_0_* from 19.9 to 21.9, i.e. by up to 10%. As for the dilution effect induced by *Q. pubescens*, the impact of canker was greater in a forest where the chestnut tree frequency was set at its lowest value, where *R_0_* rose by up to 24%, and took its minimal value, i.e. ∼2%, when the frequency of chestnut trees was at its highest. The production of native parasitoids in galls formed on *Q. pubescens* (*effect iii*) induced the strongest variations of the *D. kuriphilus R_0_* with reductions up to −66%, −31% and −7% in patches made of 16%, 60% and 91% of chestnut trees, respectively (Figure 4C). As shown by our statistical analysis, this effect is proportional to the density of *Q. pubescens*, and is then stronger in patches with low chestnut tree frequencies. Noteworthy, the calculation of those reductions of the invasive potential of *D. kuriphilus* per percent of change in the frequency of *Q. pubescens* (Figure 4D-F) and *C. parasitica* (Figure 4E) confirmed that these effects are mostly cumulative as adding a percent of pubescent oaks or canker infected chestnut trees had the same impact whatever their occurrence in the stand.

Overall, our model analysis showed that *Q. pubescens* and, at a lower extent, *C. parasitica* interferes with *D. kuriphilus* invasion and significantly reduce or increase its population growth rate through dilution of oviposition, production of native parasitoids and amplification of host detection. Furthermore, the alternative niche provided by *Q. pubescens* galls to the control agent, *T. sinensis*, could ease its establishment and spread in mixed forest. We then intended to use our modelling to predict the hyperparasitism rate of the galls formed by *A. kollari* and *A. dentimitratus* on pubescent oaks by *T. sinensis*, and the subsequent impact on *D. kuriphilus* control levels.

### *T. sinensis* parasitism of *Q. pubescens* galls slightly improve its biological control efficacy

The predicted rates of infection of *A. kollari* and *A. dentimitratus* by *T. sinensis* were shown to reach up to 1.51% and 5.49%, respectively, when considering the average estimate of *T. sinensis* searching area for each of the two parasites (Figure 5B). These predictions varied substantially when considering lower and larger values of *T. sinensis* searching areas with variations ranging from 0.39% to 4.79% for *A. kollari* and from 0.29% to 30.5% for *A. dentimitratus* (Figure 5A,C). Interestingly, their remaining variations, with respect to the chestnut tree frequency, showed a similar trend. The infestation rate of *Q. pubescens* galls by *T. sinensis* is typically null at chestnut tree frequencies lower than 40%, it increases to reach its highest level for chestnut tree frequency of 50-70%, and then decreases in stands dominated by chestnut trees, where strong oscillations of *D. kuriphilus* and *T. sinensis* abundances are typically predicted.

**Figure 5.**
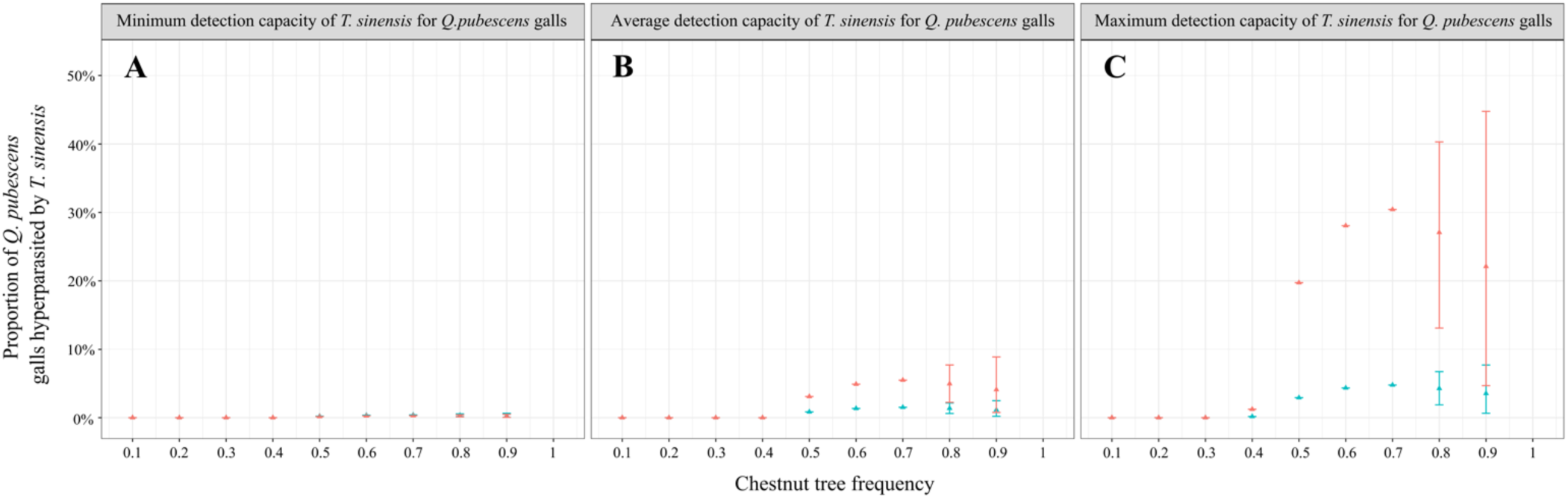
Predicted rates of hyperparasitism of *Q. pubescens* galls *by T. sinensis*. Proportions of *A. kollari* (blue) and *A. dentimitratus* (red) galls infested by *T. sinensis* were calculated for the minimum (A), average (B) and maximum value (C) of the 95% confidence interval of the searching area of the control agent (SM3). Simulations were run for chestnut tree frequency ranging from 0 to 1, and while considering all other trees as pubescent oaks.

These theoretical results suggest that infestation of *Q. pubescens* by *A. kollari* and *A. dentimitratus* could i) increase the *T. sinensis* demography by providing alternative hosts, and ii) constitute a refuge for the control agent population during periods of low *D. kuriphilus* abundance. However, the impact of these complementary effects on the *T. sinensis* control efficacy remains limited since the resulting gain in control at high frequency of *Q. pubescens* never exceed +2.2% (SM5).

## Discussion

Since its first detection in the French Eastern Pyrenees in 2013 (49), the worldwide pest *D. kuriphilus* successfully invaded and established in the corresponding natural chestnut tree populations. Across the 8 studied localities, 50-90% of chestnut trees were found infested in 2019, and, in 3 of these localities, the *D. kuriphilus’s* rates of infestation exceeded the threshold of 0.6 galls per bud that is thought to lead to drastic decrease of productivity (47). A strong decrease of *D. kuriphilus* infestation was observed in the two following years, resulting in less than 25% of infested trees with rates of infestation below the value of 0.3 galls per bud where tree yield loss are no longer expected (47). This reduction is mostly explained by the introduction of the control agent, *T. sinensis,* in 2014, which has been shown to infest about 90% of *D. kuriphilus* larvae in this studied area in 2020 (28). Such widespread and high parasitism rates of the control agent are in line with the rate at which *T. sinensis* was previously shown to spread to reach >90% rates of *D. kuriphilus* infestation within 5-7 years in Italy (60). While the decrease in pest infestation was shared in all sites, we observed significant spatial heterogeneity in the initial infestation levels as well as in their reduction rates from 2019 to 2021 (Figure 2). As mentioned in the introduction, such variations can typically result from heterogeneity in bottom-up and top-down effects affecting the spread of the invasive species. We indeed showed that *D. kuriphilus* spatial variations in infestation levels were positively correlated to the chestnut tree frequency (Table 1A,B), confirming previous studies in the French Eastern Pyrenees (28) and in other European places (61–62). This bottom-up regulation effect is then consistent with the resource concentration hypothesis (63) which states that more dense stands of a plant shall recruit more herbivores per plant unit, with a stronger effect for specialist herbivores. Meanwhile, the rates of *D. kuriphilus* infestation by native parasitoids and fungi were previously found to be both low, i.e. less than 4.5 and 6.7% respectively, and highly similar between localities, so that such top-down factors could not explain observed variations in infestation (28). In addition to the predominant bottom-up effects shown to regulate *D. kuriphilus* invasion in the French Eastern Pyrenees, we further observed that two native species, *Q. pubescens* and *C. parasitica*, have significant indirect effects on its prevalence and rate of infestation of chestnut trees.

First, we showed that the presence of *Q. pubescens*, which is phylogenetically close to the chestnut tree and the major host of most cynipid species, can disturb *D. kuriphilus* host detection and lower its invasion potential (*effect i*). The negative correlation observed between the *D. kuriphilus* rate of infestation and the pubescent oak frequency (among non-host trees) (Table 1A) indeed support such an hypothesis, and is consistent with the previous record of a lower number of galls per buds in chestnut tree stands mixed with *Q. pubescens* (61). This effect is likely explained by the waste of time spent by *D. kuriphilus* on trees that are not suitable for oviposition (18) that leads to a typical dilution effect (19). The integration of this indirect effect into our *D. kuriphilus* - *T. sinensis* dynamical model then showed that the *D. kuriphilus* invasive growth rate (*R_0_*) decreases linearly with the frequency of *Q. pubescens* among non-chestnut tree species. The magnitude of this dilution effect was further showed to be maximal in patches with low chestnut tree frequencies, while it remains limited in chestnut tree dominated forests. This suggests that the reduction of *D. kuriphilus* oviposition, and therefore of its invasion potential, is maximised in mixed forests and almost null in chestnut tree coppices. In the conditions encountered in our study area, the reduction of the pest’ *R_0_* is expected to reach a maximum of 11% in the locality ‘Ceret’ where the forest environment is composed of 19% of chestnut trees and where 26% of the non-host trees are pubescent oaks.

Second, we assessed the prevalence levels of *C. parasitica* infection in the chestnut tree populations to test if, as suggested in Germinara et al. (42), the damages induced by this parasitic fungus could enhance the release of phenolics compounds and facilitate the *D. kuriphilus* detection of canker-infected trees (*effect ii*). We indeed showed, at both sites and individuals scales, higher prevalences of *D. kuriphilus* infestation on chestnut trees infected by *C. parasitica* than on healthy hosts (Table 1B,C). These field data then suggest that canker infection could amplify the spread of *D. kuriphilus* by facilitating host detection, in line with the invasional meltdown hypothesis (64–65) stating that introduced species could promote the spread of pathogen or other invasive species if they enhance their infection success (66). Our model analysis then showed that this amplification of host detection contributes to increase the pest’ *R_0_* proportionally to canker prevalence levels, and that the magnitude of this effect is larger in mixed forest with low chestnut tree frequency while it is minimal in chestnut tree coppices (Figure 4B). Although this increase of the pest invasion potential remained lower than +13%, according to the maximum canker prevalence rate recorded in 2021, the recent rise of *C. parasitica* infection levels across the French Eastern Pyrenees (unpublished data) could enhance the intensity of this indirect amplification effect. Such an effect, combined with the predicted oscillatory dynamics between *D. kuriphilus* and its control agent (28, 41), suggest that the re-emergence of the pest could be expected in the studied area, and facilitated by this parasitic fungus.

Beyond its impact on *D. kuriphilus* oviposition, *Q. pubescens* was also shown to contribute to the production of native parasitoids able to infest the pest’ larvae (*effect iii*). While the native parasitoids *Eurytoma setigera, T. auratus, T. flavipes* and *Eupelmus sp.* were previously found in *D. kuriphilus* galls collected on chestnut trees (28) and known to be able to infest galls formed on pubescent oaks (40), only 2 of them (*T. auratus, T. flavipes)* were observed in the 208 oaks galls used in our barcoding analysis. We then showed a positive correlation between native parasitoids emergence rates from chestnut tree galls and pubescent oak abundances in each site. Altogether, these results support the assumption that chestnut tree forests mixed with *Q. pubescens* could lead to a stronger top-down control of the pest than in chestnut tree coppices (43). The integration of this empirically-supported effect of *Q. pubescens* on native parasitoids production in our modelling then provided further quantitative insights into its strong potential impact on the pest *R_0_*. We indeed predicted *D. kuriphilus R_0_*’ reductions of up to −66% when the chestnut tree frequency is minimal (0.16) and the rest of the tree community is made of pubescent oaks (Figure 4C). Although such high native hyperparasitism rates are rarely observed in chestnut tree galls, the field observations of *T. flavipes* hyperparasitism rates up to 31.75% in Italy (67) supports the potential existence of such high top-down pressures induced by native parasitoids. Meanwhile, the maximum density of pubescent oaks recorded among our sites only reaches 120 individuals per hectare, representing 26% of the non-host species community in this site. We therefore anticipate that the magnitude of this effect should remain lower than 17% among our sampling sites, in line with the observations of *D. kuriphilus* hyperparasitism rate by native parasitoids that are typically lower than 10% (40,68–69). Overall, these predictions confirm the general understanding that parasitoids naturally occurring in the invaded forest are insufficient to regulate the *D. kuriphilus* populations (50–51).

Our barcoding analysis also provided evidence that *Q. pubescens* galls are infested by the control agent *T. sinensis*, as previously observed by Ferracini et al. (44). These authors recorded *T. sinensis* emergence from 15 different gall-forming parasite species of pubescent oaks, showing the ability of the pest to switch to closely related host species. On the contrary, they did not record any switch to galls formed on *F. sylvatica* by *M. fagi*, which is confirmed by our study as no *T. sinensis* was found among the 263 galls of this parasite species that we collected on beeches. The repeated observation of a plasticity of *T. sinensis* in its host choice sharply contrasts with the expected specificity of biological control agent towards their target. In addition, the inter-specific competition occurring between native and introduced parasitoids for *D. kuriphilus* and oak parasites galls may induced a loss of 14% of native parasitoids species and 32% of their individuals (43,51,70–71). However, no long-term assessment of this (not so specialist) introduced species impact on native parasitoids species richness or abundance has been performed in oaks galls. Taken together, these observations should encourage to further evaluate the potential impact of this biological control agent on non-target endemic species. In such a context, our modelling analysis further showed that *Q. pubescens* could serve as a refuge for *T. sinensis* (*effect iv*) with 5-10% of oak galls predicted to be infected by the introduced control agent when considering its average detection capacity and the average frequency of chestnut trees in the forest environment. This provides a quantitative support to the idea that alternative hosts can indeed be targeted, and eventually increase the building of biological control agent population (24). The rate of oak galls infestation was shown to decrease at high chestnut tree frequency, where *D. kuriphilus* - *T. sinensis* typically enter oscillatory dynamics (28), although such infestation then showed large variations with pubescent oaks helping the persistence of the control agent at low abundance of *D. kuriphilus*. This suggests that the presence of *Q. pubescens* in chestnut tree dominated forests could still contribute to constrain the re-emergence of the invasive population that is expected to happen as a result of *D. kuriphilus* - *T. sinensis* oscillations (28,41). Despite of these evidence for galls found on oaks to contribute to the production and persistence of *T. sinensis*, the positive variations of *D. kuriphilus* biological control efficacy were predicted to remain lower than 2.3%, with a maximal intensity reach in mixed forest.

Overall, this integrative study provides empirical and modelling insights into the indirect effects induced by *Q. pubescens* and *C. parasitica* on the invasion and biological control of *D. kuriphilus* in the French Eastern Pyrenees. The dilution effect exerted by *Q. pubescens* on the oviposition rate of *D. kuriphilus* (*effect i*) was shown to typically dominates the amplification of chestnut trees detection induced by *C. parasitica* infection (*effect ii*). In addition, *Q. pubescens* native gall-forming parasites (*A. kollari* and *A. dentimitratus*) were found hyperparasited by both native parasitoids able to infest *D. kuriphilus,* and the introduced control agent *T. sinensis*. While the production of native parasitoids have been shown to significantly reduce the pest invasion potential, the refuge provided by these alternative host for the introduced control agent is predicted to slightly improve the efficacy of *D. kuriphilus* biological control.

## Acknowledgments

This study is set within the framework of the ‘Laboratoires d’Excellences (LABEX)’ TULIP (ANR-10-LABX-41) and of the ‘École Universitaire de Recherche (EUR)’ TULIP-GS (ANR-18-EURE-0019). This work has benefited from a PhD fellowship to JLZ (Région Occitanie) and from the project ‘Modélisation hybride d’invasions biologiques’ (MITI CNRS, PI. Gourbiere S.). This work further benefited from the modelling discussions hold in the context of the ‘PHYTOMICS’ (ANR, ANR-21-CE02-0026, PI. Gourbiere S.) and ‘ELVIRA’ (ANR, ANR-21-CE20-0041, PI. Piganeau G.) projects. The funders played no role in the study design, data collection and analysis, decision to publish, or preparation of the manuscript.

## Notes

### Competing Interest Statement

The authors have declared no competing interest.

