## Supplementary Table 1 for "An integrative field and modelling study of the bottom-up, top-down and indirect effects of native species on invasive insects and their biological control : the case of the worldwide chestnut tree pest, *Dryocosmus kuriphilus*, in the French Eastern Pyrenees"

**Supplementary Table 1.** Summary table of *D. kuriphilus*, *A. kollari* and *A. dentimitratus* infestation and prevalence rates and frequencies of *C. sativa*, *Q. pubescens* and *F. sylvatica* estimated in each of the 24 sites.

| Stations | *D. kuriphilus* | | | | | | *A. kollari* | | *A. dentimitratus* | |
|---|---|---|---|---|---|---|---|---|---|---|
| | Prevalence rate | | | Infestation rate | | | Prevalence rate | Infestation rate | Prevalence rate | Infestation rate |
| | 2019 | 2020 | 2021 | 2019 | 2020 | 2021 | 2020 | 2020 | 2020 | 2020 |
| Prats de Mollo | 0.867 | 0.173 | 0.033 | 0.022 | 0.002 | <1e10-3 | 0.373 | 0.007 | 0.255 | 0.014 |
| Site 1 | 0.78 | 0.04 | 0 | 0.014 | 0 | 0 | 0.462 | 0.017 | 0 | 0 |
| Site 2 | 0.94 | 0.26 | 0.06 | 0.029 | 0 | 0.001 | 0.524 | 0 | 0.143 | 0.029 |
| Site 3 | 0.88 | 0.22 | 0.04 | 0.024 | 0.005 | 0 | 0.118 | 0.004 | 0.588 | 0.012 |
| Saint Laurent | 0.886 | 0.473 | 0.093 | 0.112 | 0.022 | 0.005 | 0.22 | 0.001 | 0.22 | 0.019 |
| Site 1 | 0.714 | 0.02 | 0 | 0.066 | 0 | 0 | 0.143 | 0.001 | 0.286 | 0.005 |
| Site 2 | 0.98 | 0.56 | 0.12 | 0.144 | 0.035 | 0.009 | 0.375 | 0.002 | 0.063 | 0.017 |
| Site 3 | 0.96 | 0.84 | 0.16 | 0.127 | 0.033 | 0.005 | 0.091 | 0.002 | 0.364 | 0.035 |
| Arles sur Tech | 0.818 | 0.513 | 0.06 | 0.062 | 0.023 | 0.001 | 0.32 | 0.003 | 0.02 | 0 |
| Site 1 | 0.92 | 0.5 | 0.04 | 0.052 | 0.008 | 0.001 | 0.79 | 0.008 | 0.053 | 0.001 |
| Site 2 | 0.703 | 0.6 | 0.04 | 0.057 | 0.013 | 0 | 0 | 0 | 0 | 0 |
| Site 3 | 0.8 | 0.44 | 0.1 | 0.079 | 0.053 | 0.002 | 0.091 | 0.001 | 0 | 0 |
| La Bastide | 0.63 | 0.033 | 0 | 0.025 | 0.003 | 0 | 0.396 | <1e10-3 | 0.076 | 0.04 |
| Site 1 | 0.9 | 0.02 | 0 | 0.006 | 0.001 | 0 | 0.389 | 0 | 0.167 | 0.05 |
| Site 2 | 1 | 0.08 | 0 | 0.063 | 0.008 | 0 | 0.4 | 0.001 | 0.029 | 0.03 |
| Site 3 | 0.1 | 0 | 0 | 0.007 | 0 | 0 | 0 | 0 | 0 | 0 |
| Llauro | 0.793 | 0.607 | 0.113 | 0.027 | 0.007 | 0.004 | 0.471 | 0.002 | 0.098 | 0.008 |
| Site 1 | 0.96 | 0.74 | 0.06 | 0.046 | 0.005 | 0 | 0.727 | 0.001 | 0.091 | 0.01 |
| Site 2 | 0.64 | 0.44 | 0.08 | 0.033 | 0.005 | 0.006 | 0.5 | 0.002 | 0.05 | 0.006 |
| Site 3 | 0.78 | 0.64 | 0.2 | 0.01 | 0.011 | 0.005 | 0.3 | 0.002 | 0.15 | 0.008 |
| Céret | 0.807 | 0.727 | 0.193 | 0.081 | 0.04 | 0.011 | 0.25 | 0.004 | 0.039 | 0.001 |
| Site 1 | 0.9 | 1 | 0.16 | 0.132 | 0.055 | 0.014 | 0.136 | 0.001 | 0.046 | 0.002 |
| Site 2 | 0.98 | 0.96 | 0.28 | 0.081 | 0.058 | 0.018 | 0.273 | 0.004 | 0.091 | 0.002 |
| Site 3 | 0.54 | 0.22 | 0.14 | 0.018 | 0.003 | 0.001 | 0.368 | 0.005 | 0 | 0 |
| Laroque | 0.634 | 0.147 | 0.18 | 0.035 | 0.003 | 0.01 | 0.257 | 0.003 | 0.057 | 0.001 |
| Site 1 | 0.76 | 0.32 | 0.26 | 0.037 | 0.009 | 0.015 | 0 | 0 | 0 | 0 |
| Site 2 | 0.744 | 0.12 | 0.24 | 0.041 | 0.001 | 0.004 | 0.214 | 0.002 | 0.071 | 0.002 |
| Site 3 | 0.366 | 0 | 0.04 | 0.027 | 0 | 0.01 | 0.286 | 0.005 | 0.048 | 0.001 |
| La Massane | 0.492 | 0.11 | 0.025 | 0.027 | 0.001 | <1e10-3 | 0.377 | 0.011 | 0.557 | 0.012 |
| Site 1 | 0.393 | 0.074 | 0.046 | 0.011 | 0 | 0 | 0.364 | 0.01 | 0.591 | 0.014 |
| Site 2 | 0.5 | 0.16 | 0 | 0.044 | 0.002 | 0 | 0.435 | 0.021 | 0.565 | 0.014 |
| Site 3 | 0.54 | 0.08 | 0.04 | 0.028 | 0 | 0.001 | 0.313 | 0.003 | 0.5 | 0.009 |
| Total | 0.746 | 0.340 | 0.089 | 0.050 | 0.013 | 0.004 | 0.340 | 0.004 | 0.178 | 0.011 |

| Stations | *C. parasitica* Prevalence rate 2020 | Tree species frequencies | | |
|---|---|---|---|---|
| | | *C. sativa* | *Q. pubescens* | *F. sylvatica* |
| Prats de Mollo | 0.093 | 0.731 | 0.059 | 0 |
| Site 1 | 0.02 | 0.491 | 0.107 | 0 |
| Site 2 | 0.06 | 0.908 | 0.012 | 0 |
| Site 3 | 0.2 | 0.823 | 0.048 | 0 |
| Saint Laurent | 0.093 | 0.671 | 0.008 | 0 |
| Site 1 | 0 | 0.418 | 0.029 | 0 |
| Site 2 | 0.14 | 0.854 | 0 | 0 |
| Site 3 | 0.14 | 0.694 | 0 | 0 |
| Arles sur Tech | 0.333 | 0.58 | 0.031 | 0 |
| Site 1 | 0.3 | 0.869 | 0.009 | 0 |
| Site 2 | 0.38 | 0.286 | 0.075 | 0 |
| Site 3 | 0.32 | 0.652 | 0 | 0 |
| La Bastide | 0.013 | 0.504 | 0.023 | 0.035 |
| Site 1 | 0.02 | 0.473 | 0.008 | 0 |
| Site 2 | 0 | 0.424 | 0.082 | 0 |
| Site 3 | 0.02 | 0.588 | 0 | 0.092 |
| Llauro | 0.247 | 0.739 | 0.036 | 0 |
| Site 1 | 0.26 | 0.724 | 0.008 | 0 |
| Site 2 | 0.06 | 0.763 | 0.079 | 0 |
| Site 3 | 0.42 | 0.895 | 0.053 | 0 |
| Céret | 0.32 | 0.591 | 0.058 | 0 |
| Site 1 | 0 | 0.581 | 0 | 0 |
| Site 2 | 0.52 | 0.908 | 0 | 0 |
| Site 3 | 0.44 | 0.19 | 0.207 | 0 |
| Laroque | 0.273 | 0.562 | 0.034 | 0 |
| Site 1 | 0.48 | 0.735 | 0 | 0 |
| Site 2 | 0.04 | 0.583 | 0.017 | 0 |
| Site 3 | 0.3 | 0.311 | 0.1 | 0 |
| La Massane | 0.008 | 0.3 | 0.1 | 0.123 |
| Site 1 | 0 | 0.2 | 0.04 | 0.56 |
| Site 2 | 0.02 | 0.418 | 0.149 | 0.03 |
| Site 3 | 0 | 0.158 | 0.053 | 0 |
| Total | 0.176 | 0.604 | 0.037 | 0.012 |
