## Supplementary Table 2 for "An integrative field and modelling study of the bottom-up, top-down and indirect effects of native species on invasive insects and their biological control : the case of the worldwide chestnut tree pest, *Dryocosmus kuriphilus*, in the French Eastern Pyrenees"

**Supplementary Table 2**. Detection capacities of *D. kuriphilus* and correlation between *Q. pubescens* densities and chestnut tree galls infestation by the native parasitoids. The estimates of the detection capacities of *D. kuriphilus* for healthy ($a_{hc}$) and chestnut blight infected chestnut trees ($a_{ic}$), pubescent oaks ($a_u$) and other non-host tree species ($a_n$) are given with the upper and lower bounds of their 95% confidence interval. The y-intercept ($O_{NP}$) and slope ($P_{NP}$) of the linear relationship between *Q. pubescens* density and chestnut tree galls infestation by native parasitoids are given with their 95% confidence interval.

| Parameters | Low CI (95%) | Fit | Up CI (95%) |
|---|---|---|---|
| $a_{hc}$ / $a_u$ | 0.7 | 1.12 | 1.5 |
| $a_n$ | 0.32 | 0.66 | 1.1 |
| $a_{ic}$ | 1.22 | 1.6 | 1.84 |
| $O_{NP}$ | -0.014849 | 0.022287 | 0.059424 |
| $P_{NP}$ | 0.000101 | 0.000783 | 0.001464 |
