## Supplementary Table 3 for "An integrative field and modelling study of the bottom-up, top-down and indirect effects of native species on invasive insects and their biological control : the case of the worldwide chestnut tree pest, *Dryocosmus kuriphilus*, in the French Eastern Pyrenees"

**Supplementary Material 3**. Infestation rate and estimate of the searching efficiency of *T. sinensis* for *A. kollari* and *A. dentimitratus,* and their 95% confidence interval.

| Parasite species | Parameters | Low CI (95%) | Fit | Up CI (95%) |
|---|---|---|---|---|
| *A. kollari* | $Inf_{AK}$ | 0.001097 | 0.004243 | 0.013438 |
| | $\mu_s$ | 5.433129e-10 | 2.105671e-09 | 6.699653e-09 |
| *A. dentimitratus* | $Inf_{AD}$ | 0.000816 | 0.015625 | 0.095414 |
| | $\lambda_s$ | 4.04344e-10 | 7.798834e-09 | 4.965897e-08 |

To estimate the searching efficiency of *T. sinensis*, we used the only available data on the infestation rate of these oak parasites by *T. sinensis* (44), which reported 3 and 1 *T. sinensis* emergence from 707 *A. kollari* galls and 64 *A. dentimitratus* galls, respectively. We assumed that the corresponding rates of parasitism resulted from the equilibrium dynamic of our modelling, which provided the relationships;

$$\mu_S = \frac{-\ln(-Inf_{AK}+1)}{P^*}$$ Equation 6

$$\lambda_S = \frac{-\ln(-Inf_{AD}+1)}{P^*}$$ Equation 7

where $Inf_{AK}$ and $Inf_{AD}$ represent the estimated rates of infestation for *A. kollari* and *A. dentimitratus* and , P* stands for the  abundance of *T. sinensis* predicted at equilibrium. The number of *T. sinensis* at equilibrium was derived from our modelling while considering a mixed forest with 45% percent of chestnut trees, to mimic the forest stand where infestation rates were estimated (44).
