## Supplementary Table 4 for "An integrative field and modelling study of the bottom-up, top-down and indirect effects of native species on invasive insects and their biological control : the case of the worldwide chestnut tree pest, *Dryocosmus kuriphilus*, in the French Eastern Pyrenees"

**Supplementary Table 4**. Definition and estimates of the parameters of the *D. kuriphilus* host - *T. sinensis* and native parasitoids dynamical model.

| | **Symbol** | **Estimate and 95% confidence interval** | **References** |
|---|---|---|---|
| ***Castanea sativa*** | | | |
| Genetic susceptibility | $S_e$ | 0.68 | 28 |
| ***Dryocosmus kuriphilus*** | | | |
| Egg survival | $S_{He}$ | 0.535 ± 0.003 | 72 |
| Larvae survival | $S_{Hl}$ | 0.936 ± 0.01 | 73 |
| Adult survival | $S_{Ha}$ | 0.952 ± 0.01 | 74 |
| Fertility | $F_H$ | 90 ± 6 | 28 |
| Carrying capacity per hectare | k | 14 880 000 | 28 |
| ***Torymus sinensis*** | | | |
| Larvae and adult survival | $S_P$ | 0.913 ± 0.004 | 75 |
| Searching area | $a_s$ | $4.4 \times 10^{-4} \pm 0.4 \times 10^{-4}$ | 28 |
| ***Native hyperparasitic fungi*** | | | |
| Survival to fungal infection | $F_{nf}$ | 0.946 ± 0.001 | 28 |
