## Supplementary Figure 1 for "An integrative field and modelling study of the bottom-up, top-down and indirect effects of native species on invasive insects and their biological control : the case of the worldwide chestnut tree pest, *Dryocosmus kuriphilus*, in the French Eastern Pyrenees"

**Supplementary Material 5.** Variations of the *D. kuriphilus* biological control efficacy with respect to *Q. pubescens* frequency and *C. parasitica* prevalence. The impacts of *Q. pubescens* frequency (x-axis) and *C. parasitica* prevalence (y-axis) on the limitation of *D. kuriphilus* abundance by *T. sinensis* were tested for the minimum (A), average (B) and maximum (C) chestnut tree frequency observed in the study area. Variations in the level of control were expressed as a percentage of variations of the control efficacy in their absence. Positive and negative variations of the control rate of the pest population appear in blue and red, respectively.

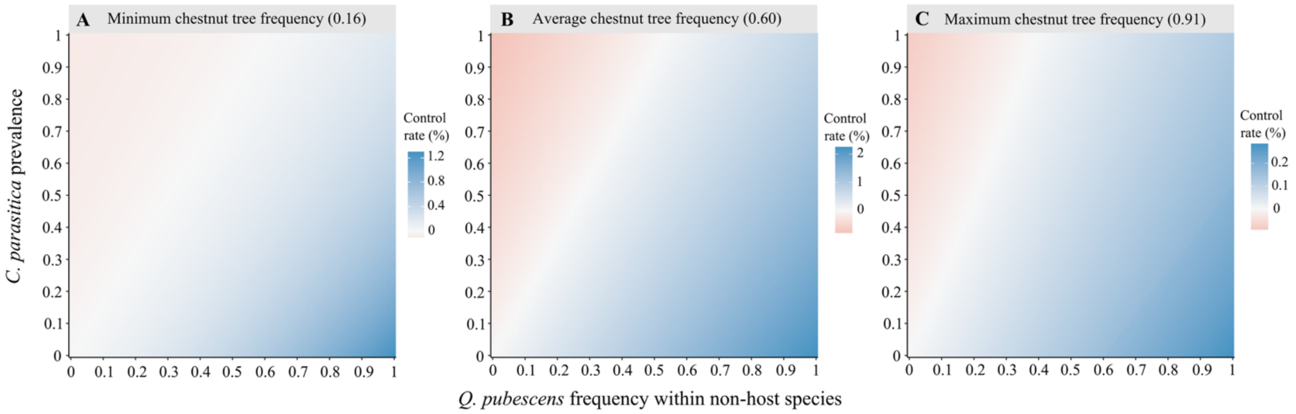
